# Methods for the Refolding of Disulfide-Rich Proteins

**DOI:** 10.1101/158162

**Authors:** Christopher N Haggarty-Weir, Roma Galloway, William Godfrey

## Abstract

*Eschericia coli* remains the workhorse producing recombinant proteins given its ease of handling, access and genetic manipulation using standard laboratory techniques. However, disulfide-rich proteins can be difficult to produce in *E. coli*, in large part due to the reducing environment of the bacterial cytoplasm. Refolding from insoluble inclusion bodies can be a viable strategy for generating substantial quantities of disulfide-rich protein. For the best chance of successfully refolding a protein, it is vital to carry out a variety of small-scale test refolds under a swathe of conditions including altering the concentration of urea, salts, reduced and oxidized glutathione, temperature, length of refold time and protein dilution factor. Once a protein has undergone refolding it is vital to determine that the final product is natively folded given the chance of soluble misfolded protein. For determination of correct folding, a variety of techniques can be employed, and ideally, numerous should be used together. For proteins that possess enzymatic function the gold standard to assess correct folding is an activity assay. Non-enzymatic proteins can be assessed using a combination of circular dichroism and nuclear magnetic resonance spectroscopy. These techniques should be utilized alongside mass spectrometry, Western blotting and SDS-PAGE.

## Introduction

The first recombinant expression of a protein in *E. coli* occurred in 1976 at Genentech, where researchers produced somatostatin (Itakura K et al, 1977), thus revolutionizing the field of biotechnology and allowing for a great leap forward in biochemistry research. Since then, more advanced systems have been developed for recombinant protein production, such as the use of yeast, insect cells, plants, mammalian cells and even cell-free systems (Bill RM, 2014). But despite these advancements, *E. coli* remains a workhorse for biochemists to make proteins with. Today up to 30% of biopharmaceuticals, 50% of commercial proteins and over 70% of proteins produced in research settings are made in *E. coli* (Bill RM, 2014).

However, the system is not without its limitations. If one requires posttranslational modifications, then a eukaryotic system would be preferred, and other issues such as incorrect disulfide bond formation and insoluble inclusion body production can present significant obstacles to the protein biochemist working with an *E. coli* expression system (these and other issues are reviewed in Rosano GL and Ceccarelli EA, 2014). Proteins that are rich in disulfides may incur difficulty in cysteine oxidation (which occurs in the periplasm) due to the reducing environment of the bacterial cytoplasm, which can lead to production of misfolded protein or inclusion body formation (Rosano GL and Ceccarelli EA, 2014).

Inclusion body formation may also occur due to the paucity of spatio-temporal control over foreign gene expression given that the recombinant protein is being manufactured in a potentially very different microenvironment (Rosano GL and Ceccarelli EA, 2014). There are numerous strategies to ameliorate these issues, including optimization of protein expression (i.e. altering concentration of the inducer and induction temperature), use of certain tags (such as MBP and MalE), using another organism, and of course, attempting to refold your desired protein from the inclusion bodies. Whilst challenging, it is can be possible to obtain sizeable and pure protein yields from inclusion bodies, and so carrying out a refolding trial should always be considered before abandoning *E. coli* as a production system (Burgess RR, 2009). This methods paper will outline one such refolding strategy that is scalable, suitable for disulfide-rich proteins, as well as downstream purification methods and the biophysical characterization of a final product. This method assumes use of a polyhistidine tag on the recombinant protein, however the general refolding methodology is transferable to various tags.

## Methodology

### Test expression

After plating freshly transformed *E. coli* containing the synthetic gene of interest onto LB agar with a suitable selective marker (i.e. kanamycin), inoculate a colony into a 50 - 100 ml flask (preferably baffled) with 10 ml of super broth plus antibiotic, growing overnight at 37 °C with agitation. The following day prepare 9 × 300 ml baffled flasks each with 30 ml of super broth plus antibiotic and inoculate 1 ml of the starter culture into each and then place in a 37 °C incubator at 220 rpm (see fig. 1 for an overall flow-chart). Grow until an optical density at 600 nm (OD_600_) of 0.6 – 0.8, then split the flasks so three will remain at 37 °C, three are to be placed in a 30 °C incubator and three are to be placed in an 18 °C incubator (for the latter two incubators, remove the flasks from the 37 °C incubator when the OD_600_ is just before 0.6 so as to pre-equilibrate the temperature before induction). Of each three lots of flasks, induce one of each with 0.25 mM, 0.5 mM and 1 mM isopropyl β-D-1-thiogalactopyranoside (IPTG) after taking a 1 ml pre-induction sample. For the flasks at 37 °C, grow for 4 hours, take a 1 ml post-induction sample and then harvest the rest by centrifugation (8 – 10 000 rpm for 10 minutes). For the flasks at 30 °C, and 18 °C, grow for 8 and 16 hours respectively before collecting a post-induction sample, measuring the final OD_600_ and harvesting the flask contents.

**Figure 1.**
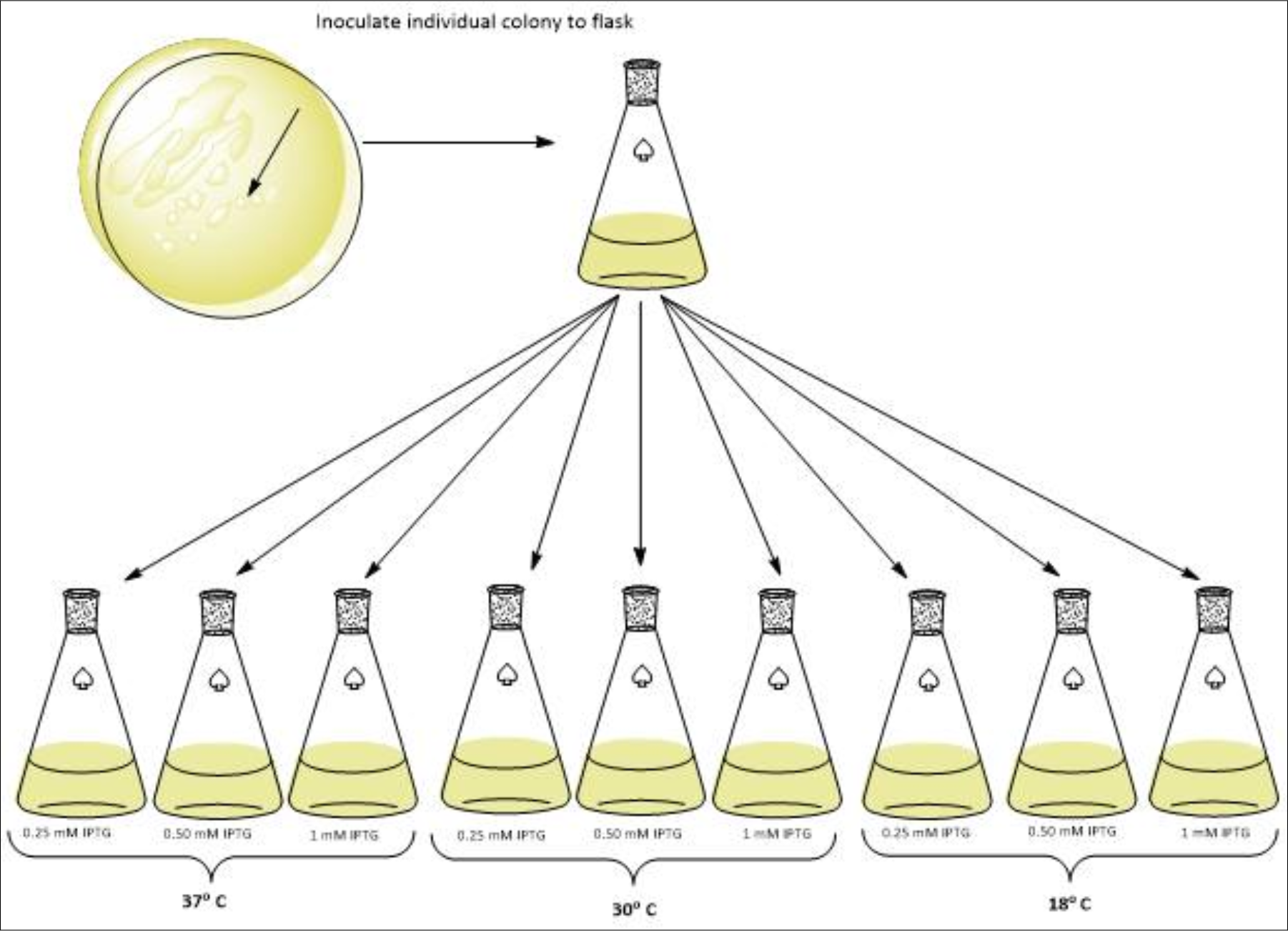
Flow-chart of Protein Test Expression: This figure presents the overall initial setup from post-transformed *E. coli* colony picking and starter culture, to optimization screening of both IPTG concentration and temperature.

Take 250 μl of each pre-induction sample, and 50 ul of post-induced sample, centrifuge, remove supernatant and add 10 μl of 1 × reducing sample buffer (RSB), then boil for 1 minute at ∼97 °C. Carry out an SDS-PAGE analysis on these samples. For each sample showing evidence of induction for recombinant protein production, centrifuge and resuspend in 2 ml lysis buffer (50 mM Tris pH 8, 250 mM NaCl, 0.1% triton X-100, 2 mM EDTA, spatula tip of lysozyme and 1.5 ul of DNasel), then process via sonication (90 sec on/100 sec off at 35% amplitude for 7.5 minutes) or mechanical cell disruption. Take a 50 ul sample post-lysis and centrifuge at around 9 000 G for 1 minute, then separate the supernatant from the pellet. Take 6 ul of the supernatant and add 10 ul of 1 × RSB; for the pellet, add 50 ul of water, resuspend thoroughly so that 6 ul can be taken and added to 10 ul of 1 × RSB. Boil samples for 1 minute at ∼97 °C. Carry out an SDS-PAGE analysis on these samples to determine that you have protein expressing in the inclusion body fraction.

### Protein production scale-up and inclusion body production

Set up 1 × 500 ml of the culture media broth with antibiotic (per construct) and inoculate each with a starter culture grown overnight from your transformed bacteria. Grow at the temperature and induce with an IPTG concentration as previously determined. Process as described in the “test expression” section (adding protease inhibitors such as PMSF or a protease inhibitor cocktail at the cell lysis step, and scaling up the volumes).

Once there is the post-lysis pellet, carry out 2 - 3 more washes with 25 mls of lysis buffer and if possible, sonicate between each wash before spinning the pellet down. Weigh out the pellet and if there is a need to store, do so at -20 °C. Solubilize the pellets with 5 ml/g of solubilization buffer (5 M guanidine HCl, 250 mM NaCl, and 20 mM Tris pH 8, and 20 mM Beta-mercaptoethanol βME). Leave this on a rotor for over an hour at room temperature then centrifuge at around 7500 G for 10 mins. Transfer supernatant to a tube with 3 mls settled Ni-NTA resin (assuming protein has a poly-His tag). Leave this on the rotor overnight at room temperature.

### Affinity purification

Let the resin settle and set up disposable tubes for the resin and sample. Add most of the fluid above the resin to the columns and collect the first 2 CVs (6 ml) as breakthrough fractions (BT) and collect the rest in a separate tube. Now mix the resin and add to the column, allowing it to run empty into the aforementioned tube. Get fresh collection tubes, add 10 CVs (30 ml) of solubilization buffer and collect as the guanidine wash fraction. Add 10 CV of 8M Urea wash/concentration buffer (8 M urea, 250 mM NaCl, 20 mM Tris pH 8 and 20 mM βME) and collect. Add 10 CVs of elute buffer with 20 mM BME and use 20 × 1.5 ml tubes to collect fractions.

Add 5 × 0.5 CV of elute buffer (8 M urea, 250 mM NaCl, 20 mM Tris pH 8, 1 M imidazole 20 mM βME), mix and leave for 15 minutes, elute and collect in tubes. Add 10 CV of solubilization buffer and wash through (collect in a tube), then add 5 ml more, mix and reapply the breakthrough (after taking samples for gels) and re-bind overnight. One may get numerous batches of protein this way.

Run samples on a gel via SDS-PAGE; either take neat samples if there is significant protein (measure Abs_280_), or carry out a TCA precipitation. Combine all fractions that have protein in them and aliquote out for the refold test.

### Refolding trials

Take 1 ml of your eluted protein sample and add 100 ul of 200 mM DTT (made in solubilization buffer without βME) and call this your reducing sample (R). Take 1 ml of another aliquote and add 100 ul solubilization buffer to make your Non-Reducing sample (NR). Leave at room temperature for 2 hours.

In a tube, prepare 4 mls of 0.1 M reduced glutathione, and in another make up 500 ul of 0.1 M oxidized glutathione. Get 8 × 50 ml tubes and label half R and half NR. Of the R half, label each one as follows (then do the same for the NR tubes)- 0 M Urea, 1 M Urea, 2 M Urea, 3 M Urea. These tubes are to hold 20 mls of solution by the end of the set up.

Add urea salts to the tubes (0 g for the 0 M tubes, 1.2012 g for the 1 M tubes, 2.4024 g for the 2 M tubes and 3.6036 g for the 3 M tubes). Next, add 200 ul of the reduced glutathione solution to each, then 20 mM Tris pH 8 (pH depends on the isoelectric point of the protein; the proteins used in this paper were all around was 5.25 - 5.75) and 100 mM NaCl. Top the tubes up to 20 mls with purified water and adjust pH. See fig. 2 for an overall schematic of the steps up to this point.

**Figure 2.**
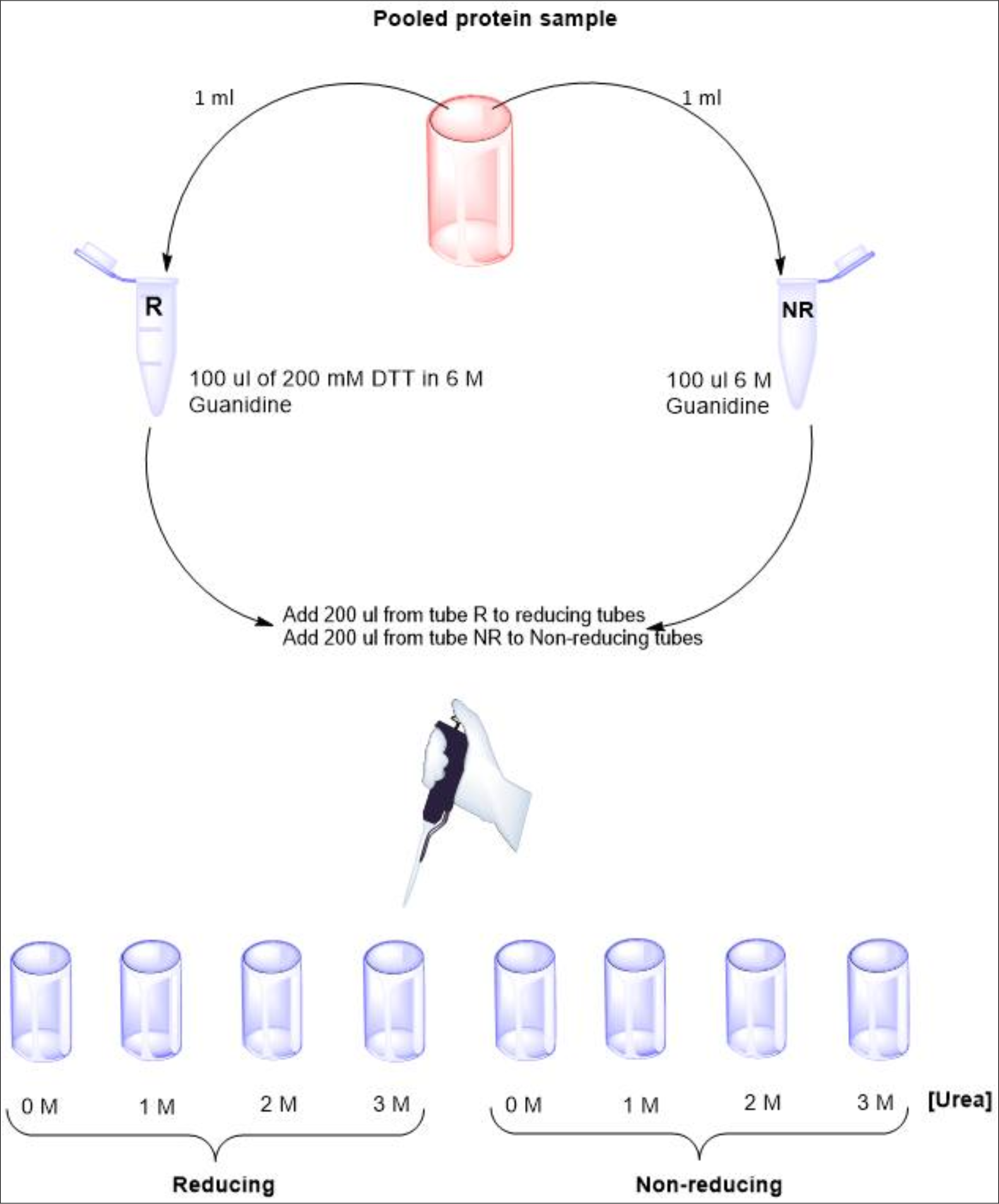
Schematic of Protein Refolding Set-up: The pool of Ni-affinity purified fractions of protein should be used for the optimization set-up of the various refolding conditions (R or NR and concentration of Urea). Slowly add 200 ul of the protein to each tube in a slow drop-by-drop fashion. Add 200 ul of the oxidized glutathione solution to each tube. Take a 2 ml sample from each tube and carry out a TCA precipitation; this will be the t_0_ sample and there should be enough to run an R and NR sample next to each other on a gel.

Leave the tubes mixing at room temperature. Each day, for 3 days, take a 2 ml sample from each tube, carry out a TCA and analyse R and NR samples. By the end of these trials you will see if the addition of urea is necessary. Further you will look for the conditions that give you the best collapse of bands over time. Note; you may not need to do a TCA if you have a great deal of protein. It is worth taking 30 ul samples and running them neat on reducing gels. Once you have the above conditions worked out, then you need to optimize the levels of glutathione used. Try the following - 1 mM R glut/1 mM Ox glut, 1 mM R glut/ 0.5 mM Ox glut, and 1 mM R glut/ 0.25 mM Ox glut.

By now the conditions for scaling up the refold should be clear. The final major step that can be optimized here is the dilution factor (i.e. how much protein goes into how much refold buffer). It is good to start off with 1:100, but also try 1:50 and 1:25. Also, generally it is good not to go too large with the bottle size. Only use 250 ml bottles with 250 ml of buffer (don’t use 1 L bottles with 1 L buffer unless you want to attempt to optimize this).

### Scale up

Note 1. This assumes a 1:25 dilution factor so if using 1:100 just alter the amount of protein being added to the mix.

Note 2. The protein used for the example given below is a 16.5 kDa protein with 20 cysteine residues that form disulfide bonds, has a pi of around 5.2 and consists of two epidermal growth factor-like (EGF) domains which contain around 17% beta-sheet.

Make up a suitable quantity of 20 mM Tris/100 mM Nacl buffer, filter then chill on ice whilst bubbling through nitrogen gas for 20 mins. Get clean 250 ml bottles (the amount depends on how much protein you plan to make, often it is good only to start with 2 bottles until you have successfully made 1 batch) each with clean magnetic stirrers in them. Fill with 250 ml of the aforementioned Tris/Nacl buffer, stir in the cold room and add a protease inhibitor tablet into each.

Add 100^th^ of the volume (2.5 mls) of the stock 0.1 M reduced glutathione (if this is the concentration determined to be optimal) to each bottle. Next, add 10 mls of your protein to each bottle (for a first attempt, try 2.5 mls) in a very slow, dropwise fashion, whilst the magnetic stirrer is on. Finally, add 2.5 mls of stock oxidized glutathione (concentration will depend on what you previously worked out during optimization). Aerate the top of each bottle with nitrogen gas for 1 min, put cap on and parafilm. Leave stirring at room temp for 3 days (or the amount of days optimized before; additionally try carrying the refold out at 4 °C, or without the magnetic stirrer). Take R and NR samples each day to track the refold.

### Dialysis

After taking pre-dialysis samples to run R and NR SDS-PAGE gels of, put 2 × 250 ml refold into dialysis tubes of a suitable molecular weight cut-off and then these into a 5000 ml beaker with 20 mM Tris pH 8 and 20 mM NaCl, with two buffer changes a day for 2 days. Carry out the dialysis at 4 °C and place a magnetic stirrer in the bottom of the large beaker.

Take samples out of dialysis bags, centrifuge at top speed to remove particulates (filtering can work, but beware if the protein non-specifically binds the filter). Now take post-dialysis samples for SDS-PAGE.

### Chromatography and biophysical characterization

The final step in the purification is to obtain a purified protein sample for biophysical characterization to assess the folding of the protein (i.e. nuclear magnetic resonance, circular dichroism, mass spectrometry, and/or Western blotting) or an enzymatic activity assay if suitable for your protein. Disulfide mapping can also be extremely informative and an example of a methodology is provided by Hodder AN et al., 1996. This is beyond the scope of this methods paper, however in general, one can first utilize ion exchange followed by size exclusion chromatography to obtain highly pure samples. An example of how the sample should look at key stages in the purification process if shown in fig. 3.

**Figure 3.**
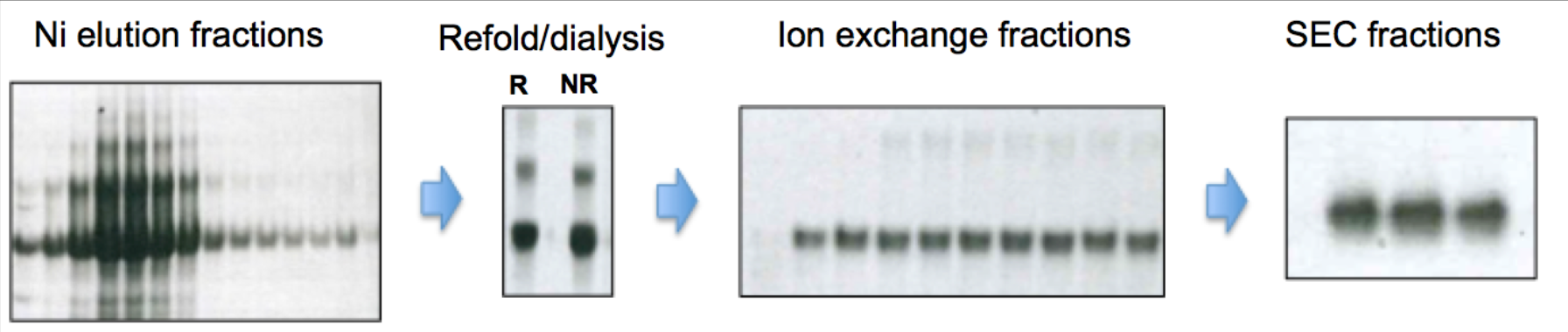
Protein purification steps: The Ni-affinity fractions eluted from the column can be run under R conditions on SDS-PAGE where both an analysis of the size of the protein can be assessed (shown as the predominant band), in addition to the higher-order species. During refolding the disulfides should shuffle and reorganize into lower order species, but generally will still include monomer, dimer and trimer. Ion exchange chromatography can be used to obtain monomer and dimer, however a final purification step may be needed to get pure monomeric protein (here demonstrated by post-SEC samples).

NMR analysis should ideally be carried out on at least a 600 MHz spectrometer for quality resolution, especially if using a non-isotopically labeled protein sample. Depending on the sample tube type, it is recommended to use around 500 μl of sample at a concentration of over 0.25 mM to reduce data collection time. Conditions such as pH, temperature and buffer composition should also be explored to optimize the quality of the spectra obtained; low salt (<75 mM), high temperature (around 37 °C) and an acidic pH generally yield the best quality spectra if your protein can tolerate these conditions. An example of NMR analysis is given in figs.4 - 6.

**Figure 4.**
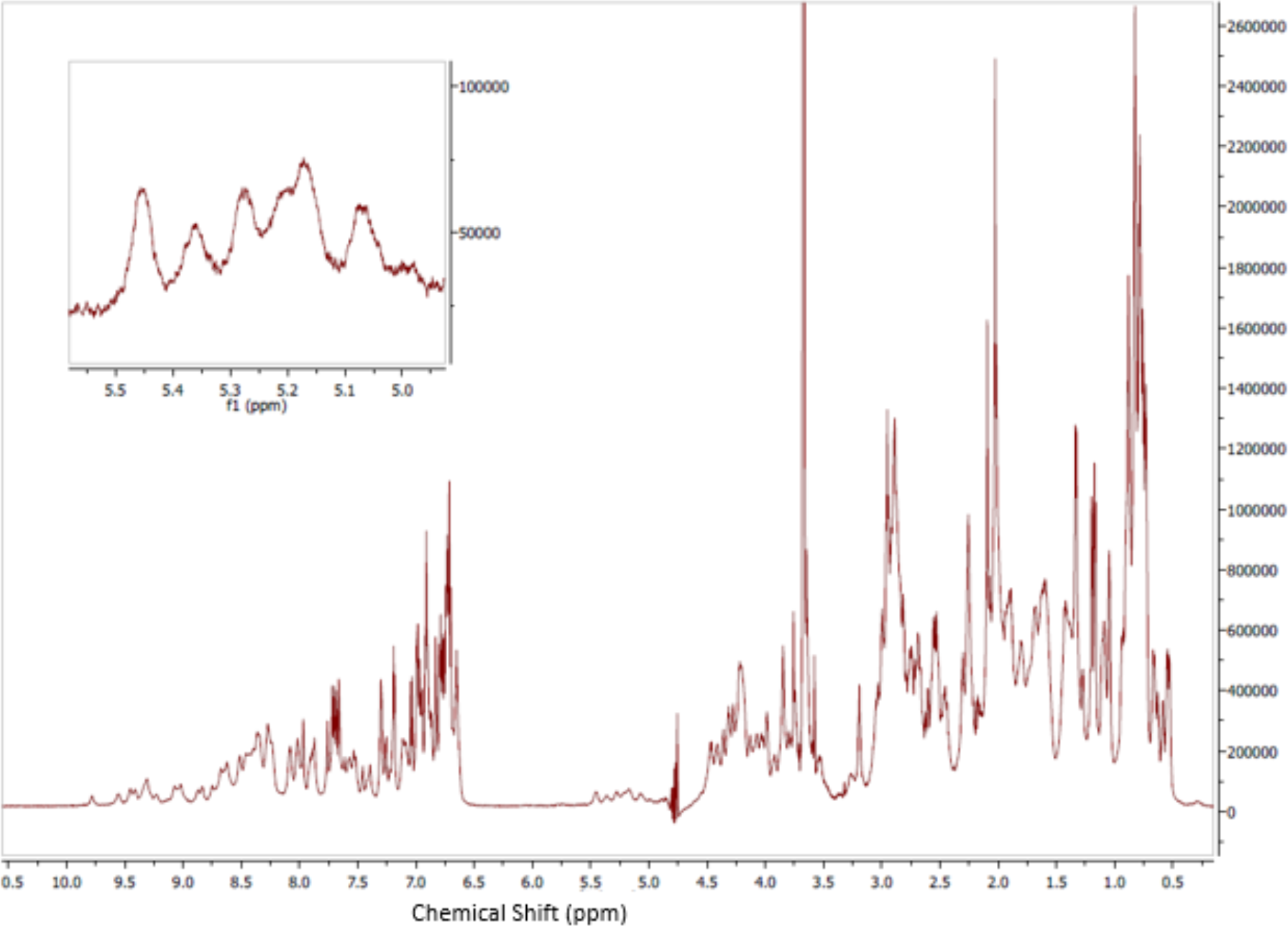
Shaka ^1^H NMR spectrum for a 2-module EGF-like protein: Data acquired from 64 scans on a Bruker 800 MHz spectrometer. Sample (280 μM) was in 40 mM PBS, 2mM Tris, 10% D_2_O, at pH 7.5 with experiment carried out at 25 °C. Detail shows α-H peaks confirming the presence of β-sheet in the secondary structure.

**Figure 5.**
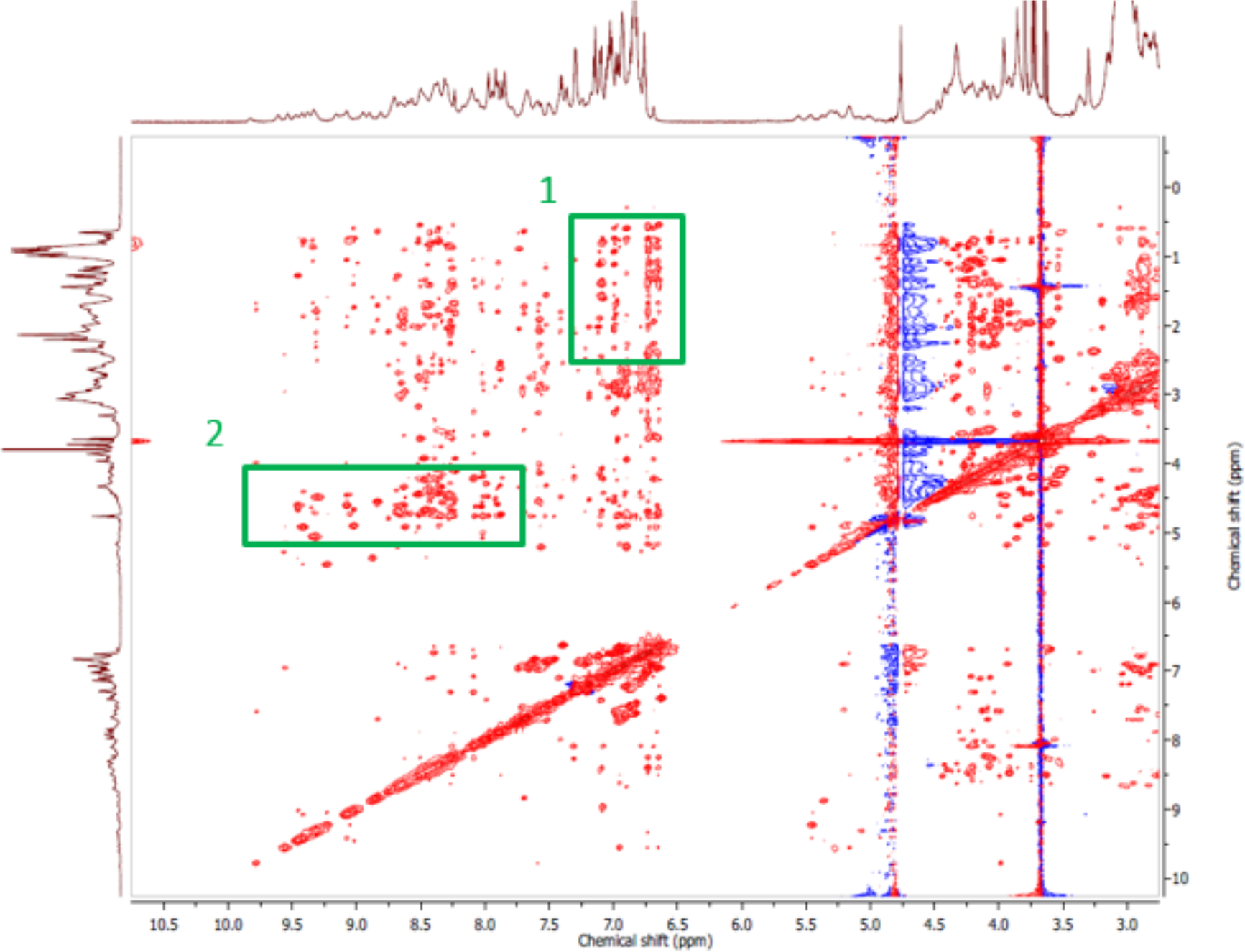
^1^H^1^H NOESY NMR spectrum for a 2-module EGF-like protein: Sample (280 μM) was in 40 mM PBS, 2mM Tris, 10% D_2_O, at pH 7.5. The experiment was carried out at 25 °C. Highlighted peaks correspond to: 1) long range methyl-H and aromatic-H interactions and 2) Backbone amide-H and α-H interactions through space. Data acquired on a 800 MHz Bruker NMR.

**Figure 6.**
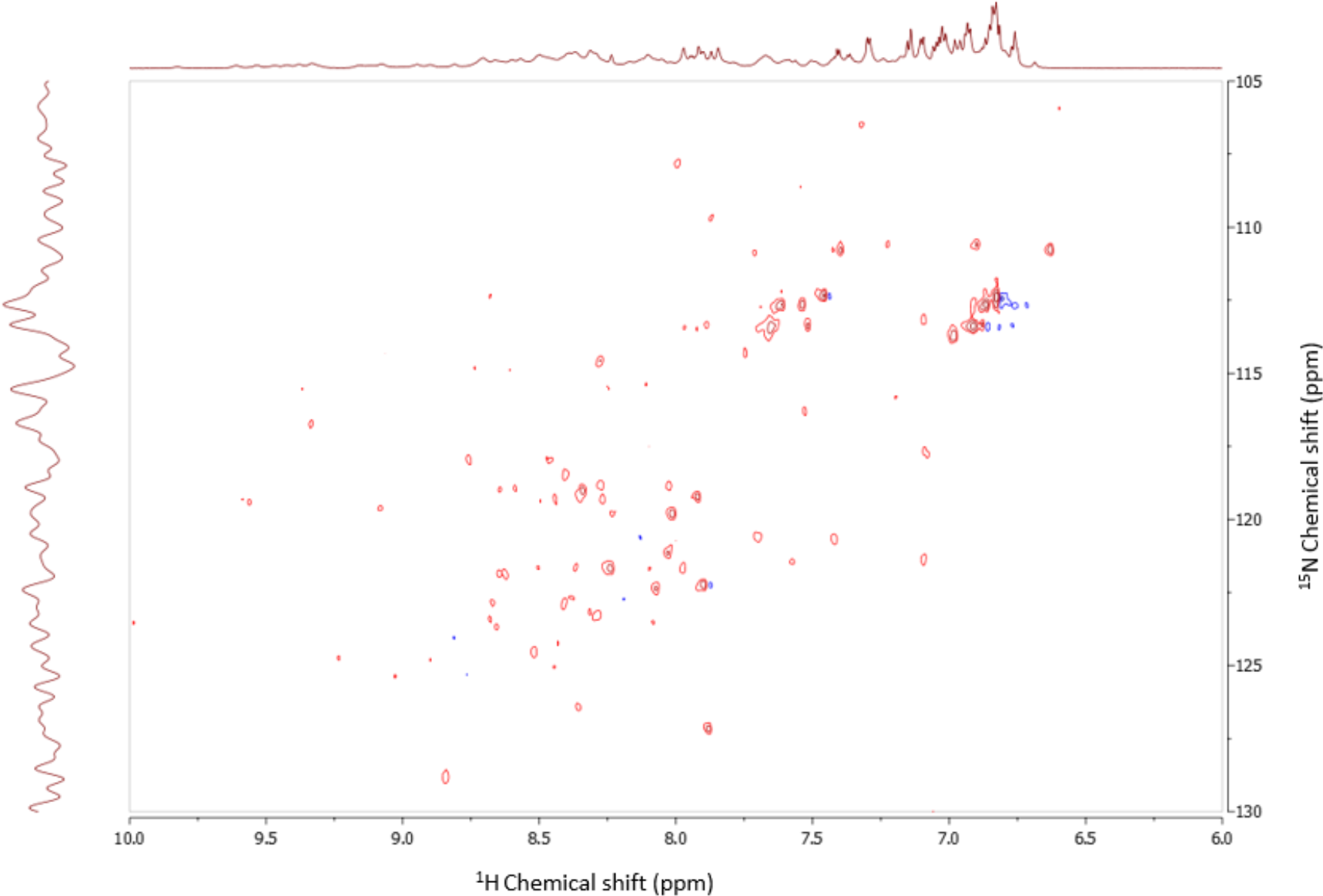
Natural abundance ^15^N^1^H HSQC NMR spectrum for a 2-module EGF-like protein: Data in red were acquired over 40 hours on an 800 MHz Bruker NMR spectrometer. The sample (280 μM) was in 40 mM PBS, 2mM Tris, 10% D_2_O, at pH 7.5. Dispersed cross peaks indicate that aggregation has not occurred. The experiment was carried out at 25 °C.

### Final Comments

This method, whilst not all-inclusive of potential refolding methods, should provide the biochemist with a reasonable level of options for trialing the refolding of their inclusion bodies. Many other conditions can be attempted, including the temperature that the refold is occurring at, or the use of L-arginine buffers. Of course, if one experiences little progress in attempting refolding, they could contact the authors for further options or move to another expression system, such as baculovirus or *Pichia*.

## Acknowledgements

All authors would like to thank the people who contributed to the Pozible malaria project run by the Walter and Eliza Hall Institute, which contributed to the work shown here. CNHW would like to thank Dr. Anthony N Hodder for the training he received in biochemistry, and the staff of Andrew Usher & Co. in Edinburgh for their service whilst this manuscript was being prepared.

## References

Bill RM. Playing catch-up with Escherichia coli: using yeast to increase success rates in recombinant protein production experiments. Frontiers in Microbiology. 2014 Mar 5;(85). http://iournal.frontiersin.org/article/10.3389/fmicb.2014.00085/full

Burgess RR. Refolding solubilized inclusion body proteins. Methods in Enzymology 2009, 463;259-282. http://www.ncbi.nlm.nih.goV/pubmed/19892177

Hodder AN, et al. The disulfide bond structure of Plasmodium apical membrane antigen-1. Journal of Biological Chemistry 1996 Nov 15;271(46). http://www.jbc.org/content/271/46/29446.abstract

Itakura K, Hirose T, Crea R, Riggs AD, Heynecker HL, Bolivar F et al. Expression in eschericia coli of a chemically synthesized gene for the hormone somatostatin. Science. 1977 Dec 9;198(4321):1056-1063. http://science.sciencemag.org/content/198/4321/1056.long

Rosano GL and Ceccarelli EA. Recombinant protein expression in Escherichia coli: advances and challenges. Frontiers in Microbiology 2014 Apr 17; 5;(172). http://iournal.frontiersin.org/article/10.3389/fmicb.2014.00172/full

